# Simultaneous CUT&Tag profiling of the accessible and silenced regulome in single cells

**DOI:** 10.1101/2021.12.19.473377

**Authors:** Derek H. Janssens, Dominik J. Otto, Michael P. Meers, Manu Setty, Kami Ahmad, Steven Henikoff

## Abstract

Cleavage Under Targets & Tagmentation (CUT&Tag) is an antibody-directed transposase tethering strategy for *in situ* chromatin profiling in small samples and single cells. We describe a modified CUT&Tag protocol using a mixture of an antibody to the initiation form of RNA Polymerase II (Pol2 Serine-5 phosphate) and an antibody to repressive Polycomb domains (H3K27me3) followed by computational signal deconvolution to produce high-resolution maps of both the active and repressive regulomes in single cells. The ability to seamlessly map active promoters, enhancers and repressive regulatory elements using a single workflow provides a complete regulome profiling strategy suitable for high-throughput single-cell platforms.

## Introduction

Throughout development, cells progress through a highly ordered series of cell fate transitions that gradually refine their cellular identities and direct their functional specializations [1]. This “epigenetic” programming is controlled by gene expression networks that tune the production of RNA transcripts from the genome. In the transcriptionally repressed state, developmental genes display a characteristic broad distribution of the Polycomb Repressive Complexes-1 and -2 (PRC-1, and PRC-2), where PRC-2 tri-methylates histone H3 Lysine-27 (H3K27me3) that extends from upstream of the transcriptional start site (TSS) out across the gene body and beyond [2]. During gene activation, cell type specific gene regulatory networks stimulate recruitment and firing of the RNA Polymerase II (Pol2) machinery and drive increased protein turnover and accessibility over transcriptional start sites (TSSs) and other cis-regulatory DNA elements that modulate gene expression (enhancers). During gene activation PRC-1 and PRC-2 are locally displaced and the H3K27me3 mark is lost. Defects in this interplay between active and repressive chromatin regulation underly a wide variety of human pathologies. However, because primary samples include complex mixtures of cells along various developmental trajectories, technologies that achieve single cell resolution are generally necessary to interrogate the molecular mechanisms that control gene expression in the normal and diseased states.

Single-cell genomic technologies that profile mRNAs (RNA-seq) or chromatin accessibility (ATAC-seq) can resolve the unique gene expression signatures and active regulatory features of distinct cell types from heterogenous samples [3]. For single-cell profiling of the repressive chromatin landscape, we have applied single-cell H3K27me3 CUT&Tag, in which an antibody that targets H3K27me3 tethers a Protein A-Tn5 (pA-Tn5) fusion protein transposome complex to chromatin [4]. To overcome the limitation of sparse or incomplete cellular profiles inherent to single cell genomics, droplet-based and nanowell platforms and combinatorial barcoding strategies dramatically increase the number of cells profiled in a single experiment [5–7]. These sparse single cell profiles can then be grouped according to shared features to assemble more complete aggregate profiles of each cell type. Platforms that simplify the workflows and data analysis have greatly facilitated profiling the gene expression signatures and active and repressive chromatin landscapes of single cells [8].

To maximize genomic information from each single cell, several methods have been developed that simultaneously profile two or more modalities, such as accessible chromatin and mRNA [9] or histone modifications and mRNA [7]. Multimodal single-cell profiling can resolve cell types that may be highly similar in the readout of one assay but show characteristic differences in the other and also allow direct comparisons between gene expression and components of the regulatory landscape in individual cells. Methods that simultaneously profile both the active and repressive epigenome could provide a more comprehensive understanding of cell fate regulation than can be obtained by profiling the active or repressive chromatin landscapes in isolation. However, multimodal methods require complex workflows and present data integration challenges, and there are no published methods that simultaneously profile the active and repressive chromatin landscape using a single workflow and readout modality.

Previously, we introduced a modified version of CUT&Tag where pA-Tn5 or Protein A/G-Tn5 (pAG-Tn5) is tethered near active TSSs and enhancers and tagmentation is performed under low salt conditions (referred to as CUTAC) [10, 11]. Low-salt tagmentation results in highly specific integration of tethered Tn5s within narrow accessible site windows to release chromatin fragments from active regulatory elements across the genome. Here we extend CUTAC to simultaneously profile regions of active and repressive chromatin within single cells by simply mixing antibodies that target both the initiating form of RNA Polymerase II and H3K27me3 followed by *in silico* deconvolution of the two epitopes. Our deconvolution strategy leverages both the different tagmentation densities and the different fragment sizes to separate active and repressive chromatin regions directly from the data without reference to external information. In this way, CUT&Tag2for1 profiles both chromatin states using a single sequencing readout. As the workflow is practically identical to that of standard CUT&Tag, we expect the method will be readily adopted for platforms already engineered for single-cell CUT&Tag.

## Results

### Pol2S5p-CUTAC maps accessibility of promoters and functional enhancers

In CUTAC chromatin accessibility mapping, pA-Tn5 is tethered to active TSSs and enhancers using antibodies targeting either Histone 3 Lysine-4 dimethylation (H3K4me2) or trimethylation (H3K4me3) [10]. We therefore reasoned that directly tethering pA-Tn5 to the initiating form of Pol2 (Pol2S5p), which is paused just downstream of the promoter, might also tagment accessible DNA under CUTAC conditions. Indeed, we found that Pol2S5p CUTAC profiles display similar enrichment to H3K4me2 CUTAC at a variety of accessibility-associated features, including annotated promoters (Fig. 1a, left) and STARR-seq functional enhancers (Fig. 1b, left) in K562 Chronic Myelogenous Leukemia cells. Pol2S5p CUTAC yielded profiles with sharp peak definition and low backgrounds relative to high-quality ATAC-seq profiles (Additional file 1a). Genome-wide, we observed high sensitivity and excellent signal-to-noise for Pol2S5p CUTAC, with more peaks called and higher Fraction of Reads in Peaks (FRiP) scores [12] when plotted as a function of fragment number (Fig. 1c). Notably, restricting CUTAC fragments to those shorter than 120 bp further improved the resolution of accessible features (Fig. 1a-b right), consistent with efficient Tn5 footprinting in exposed DNA [13]. This interpretation is supported by aligning reads from PRO-seq, a transcriptional run-on method that precisely maps the position of the Pol2 active site (Fig. 1d) [14], which shows it to be centered on average ~130 bp from the accessibility footprints genome-wide (Fig. 1e).

**Figure 1:**
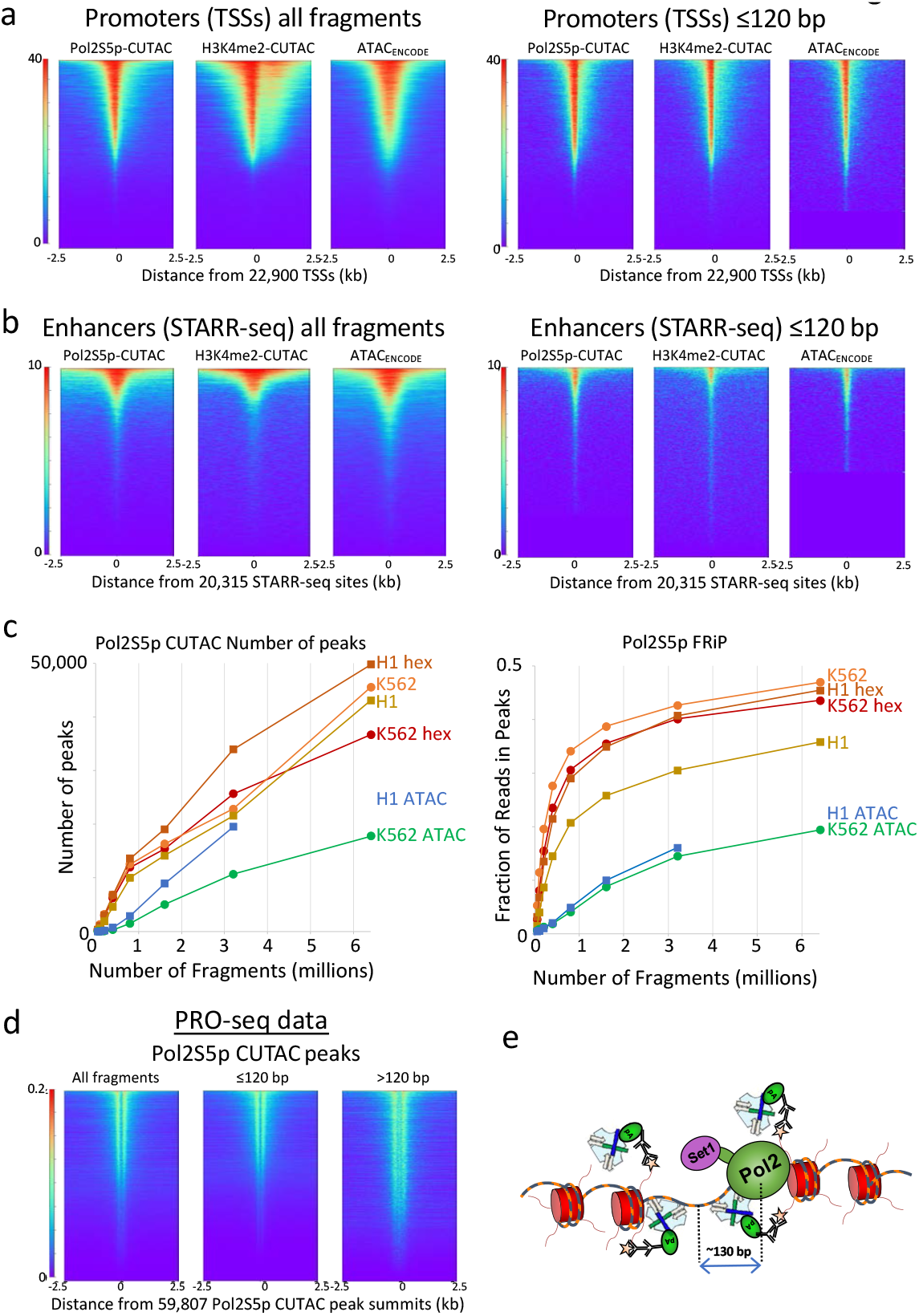
Pol2S5p CUTAC maps promoters and functional enhancers adjacent to RNAP2 genome-wide: (**a**) Heatmaps showing occupancies of Pol2S5p CUTAC, H3K4me2 CUTAC and ATAC-seq signals over promoters in K562 cells, which become sharper for subnucleosomal (1-120 bp) fragments. (**b**) Pol2S5p CUTAC, H3K4me2 CUTAC and ATAC-seq signals precisely mark functional enhancers when aligned to STARR-seq peaks. (**c**) To evaluate the data quality of Pol2S5p CUTAC, random samples of mapped fragments were drawn, mitochondrial reads were removed and MACS2 was used to call (narrow) peaks. The number of peaks called for each sample (left) is a measure of sensitivity and the fraction of reads in peaks (FRiP, right) is a measure of specificity calculated for each sampling in a doubling series from 50,000 to 6.4 million fragments. For comparison, an ENCODE ATAC-seq sample was used for K562 cells and a published ATAC-seq sample from our lab (GSE128499) was used for H1 cells. Hex samples were tagmented in the presence of 10% 1,6-hexanediol. (**d**) Run-on transcription initiates from most sites corresponding to RNAP2S5p CUTAC peaks, where PRO-seq maps the RNA base at the active site of paused Pol2 [14]. Both plus and minus strand PRO-seq datasets downloaded from GEO (GSM3452725) were pooled and aligned over peaks called by MACS2 using 3.2 million RNAP2S5p CUTAC fragments. (**e**) Model for RNAP2S5p-tethered tagmentation of adjacent accessible DNA, where the Set1 H3K4 methyltransferase di- and tri-methylates nucleosomes near stalled Pol2.

To quantify the degree of overlap between CUTAC and ATAC-seq, we called peaks from one replicate of ENCODE ATAC-seq data and aligned samples of 3.2 million fragments from Pol2S5p-CUTAC and K4me2-CUTAC data and from the other ATAC-ENCODE replicate. Based on these heat maps, we find that 98-99% of Pol2S5p-CUTAC and H3K4me2-CUTAC ≤120 bp fragments are centered over ENCODE ATAC-seq peaks, whereas the ENCODE ATAC-seq replicate shows 93% overlap (Additional file 2a-e). We attribute the better overlap of CUTAC to the ENCODE ATAC-seq standard than a replicate of the same standard to the better resolution of CUTAC over ATAC-seq, with much sharper peaks and better signal-to-noise (Additional file 2d), so that much more of the CUTAC signal is present in ≤120 bp fragments and is better centered over the midpoint of each accessible site. This close correspondence is confirmed by correlation analysis (Additional file 2e), providing direct evidence for the involvement of Pol2 in driving H3K4 methylation and chromatin accessibility [10, 15]. The fact that most promoters and STARR-seq enhancers are immediately adjacent to the paused initiating form of Pol2 is consistent with the suggestion that enhancers and promoters share the same chromatin configuration [16].

### CUT&Tag2for1 distinguishes active versus repressed chromatin based on fragment size

In comparison to H3K27me3 CUT&Tag profiles of the developmentally repressed chromatin landscape, we noticed that the CUTAC profiles include a much larger proportion of fragments that are <120 bp in both K562 and H1 embryonic stem cells (Additional file 1b). No consistent changes in fragment sizes were seen when 3-12 rounds of linear amplifications preceded PCR to minimize the competitive advantage of small fragments during the short PCR cycles used for CUT&Tag. However, including the polar organic compound 1,6-hexanediol during tagmentation resulted in a smaller fragment size distribution (Additional file 1c), with H1 cells showing a more marked effect than K562 cells, consistent with our previous finding that this increases penetrability of pAG-Tn5 [10] and with “hyperdynamic” chromatin characteristic of embryonic stem cells [17]. We reasoned that this difference in fragment size might provide a general strategy to separate active and repressed chromatin profiles using a single sequencing readout from the same cells. Accordingly, we mixed Pol2S5p and H3K27me3 antibodies and followed the CUT&Tag protocol for K562 and H1 samples with tagmentation under low-salt CUTAC conditions (Fig. 2a). We found that when compared to individual CUTAC and H3K27me3 CUT&Tag profiles, features from both targets were well-represented in CUT&Tag2for1 profiles (Fig. 2b). We applied a two-component Gaussian Mixture Model to the distribution of fragment size averages using an Expectation Maximization algorithm [18] to separate peaks into inferred Pol2S5p-CUTAC (small fragment average) and H3K27me3 (large fragment average) profiles from the mixture. We found that H3K27me3 CUT&Tag and CUTAC map nearly exclusively to their fragment size-inferred peak sets (Fig. 2c), supporting the use of fragment size as an accurate feature classifier of CUT&Tag2for1 data. These data suggest that active and repressive chromatin features can be deconvolved in a joint assay with minimal additional effort.

**Figure 2:**
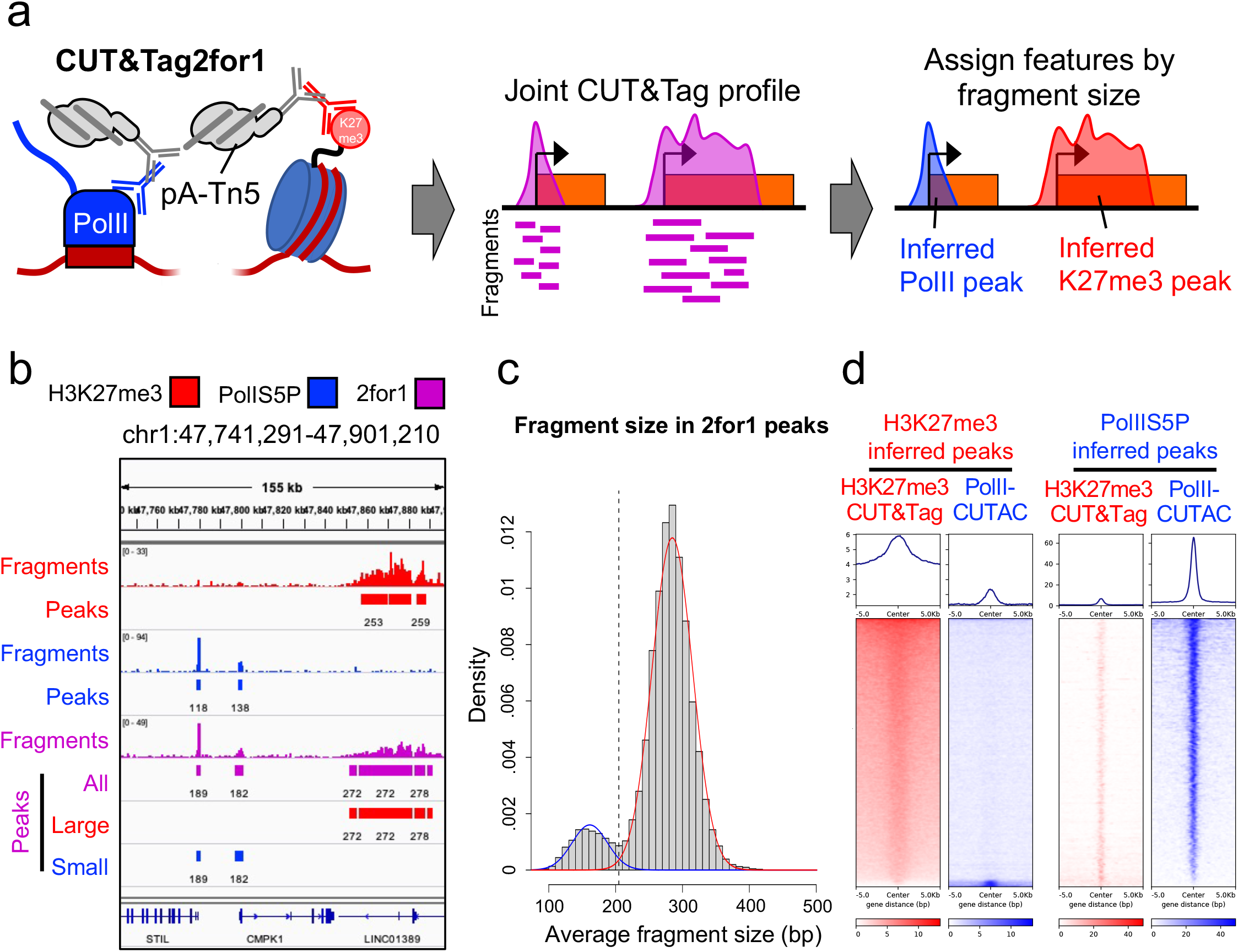
CUT&Tag2for1 simultaneously profiles active and repressive elements. (**a**) Schematic describing the CUT&Tag2for1 rationale: A joint CUTAC-CUT&Tag profile is generated using antibodies against Pol2S5p and H3K27me3, and enriched regions are assigned post-hoc based on fragment size. (**b**) Genome browser screenshot showing a CUT&Tag2for1 profile (green) in comparison with H3K27me3 CUT&Tag (red) and Pol2S5p-CUTAC (blue) for a representative region in K562 cells, along with enriched peaks. CUT&Tag2for1 large (red) and small (blue) fragment peaks as assigned by fragment size are shown. (**c**) Heatmaps describing H3K27me3 CUT&Tag (red) and RNA Pol2S5p-CUTAC (blue) enrichment at large fragment (left) or small fragment (right) peaks as defined from CUT&Tag2for1 profiles.

### CUT&Tag2for1 for single cells

Given the successful adaptation of CUT&Tag for single cell profiling [4–6] we next asked whether CUT&Tag2for1 could be adapted for single-cell chromatin characterization. We performed CUT&Tag2for1 in K562 and H1 cells, isolated single cells on a Takara ICELL8 microfluidic device, and then amplified tagmented DNA with cell-specific barcodes (Fig. 3a). Because the fragment size distributions of the two targets can exhibit considerable overlap (Additional file 1b-c), we reasoned that deconvolution can be further enhanced by considering dependencies between positionally close adapter integration sites in the genome. *i*.*e*., observation of many cut sites from a particular target makes it more likely that an integration close to this set was induced from the same target feature. In addition, we also tested whether the differences in feature and peak width for the two epitopes (Pol2S5p peaks are narrow and sharp; H3K27me3 peaks are broad and diffuse) can also help the deconvolution. By applying Bayesian statistics to model the CUT&Tag2for1 signal as a mixture of Pol2S5p and H3K27me3, we found that indeed a model that incorporates length distributions, positional dependencies, and feature widths of the two targets outperforms any of the individual parameters, and we named this novel deconvolution approach 2for1separator (Fig. 3b, Methods, Additional file 3). The fragment length distribution is encoded as a mixture of log-normal distributions over the characteristic modes of chromatin data, and the neighborhood information *i*.*e*., positional dependencies and feature widths are modeled using a Gaussian process (Fig. 3b). We then used the deconvolved signals as inputs to a peak-calling procedure to identify Pol2S5p and H3K27me3 peaks from CUT&Tag2for1 data.

**Figure 3:**
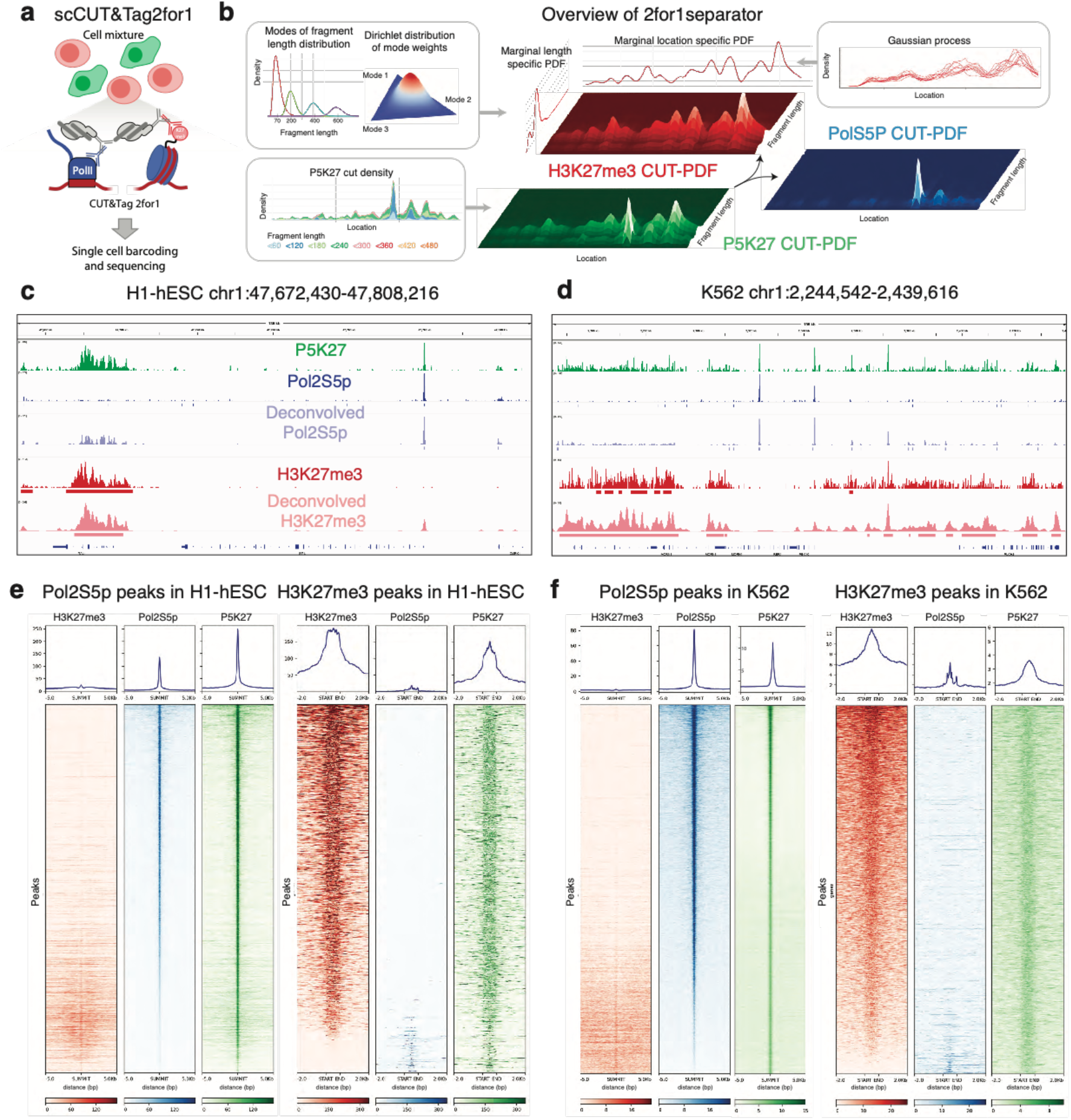
Deconvolution of CUT&Tag2for1 using fragment size and feature width: (**a**) Schematic of the single-cell CUT&Tag2for1 experimental rationale, in which two cell types are profiled in bulk in parallel and then arrayed on an ICELL8 microfluidic chip for cell-specific barcoding via amplification and mixing before sequencing. (**b**) Schematic of the deconvolution approach using a Bayesian model by considering differences in fragment length distributions and feature widths of the two targets. PDF: Probability density function. (**c**) Genome browser screenshot showing a CUT&Tag2for1 profile (green) in comparison with H3K27me3 CUT&Tag (red) and Pol2S5p-CUTAC (blue) for a representative region in H1 human embryonic stem cells (hESC), along with inferred peaks from single-cell CUT&Tag2for1 data. (**d**) Same as (c) for K562 cells. (**e-f**) Single antibody and CUT&Tag2for1 data at the inferred Pol2S5p (left) and H3K27me3 peaks (right) for H1 and K562 cells, where misclassified peak numbers and percentages are: H1 Pol2S5p (1261 = 8.2%), H1 H3K27me3 (161 = 11.0%), K562 Pol2S5p (396 = 1.0%), and K562 H3K27me3 (496 = 3.7%). In c-f, CUT&Tag2for1 P5K27 data represent the pseudo-bulk aggregate for all cells derived by pooling single-cell data, and Pol2S5p and H3K27me3 data are from single antibody data. Results were obtained by pooling cells from two single-cell replicates.

To deconvolve the Pol2S5p and H3K27me3 features from our single cell data we first created a pseudo-bulk profile by aggregating the unique reads from individual H1 and K562 cells. Our 2for1separator algorithm accurately determined Pol2S5p and H3K27me3 peaks, showing strong enrichment of the correct single antibody signals in the respective peaks (Fig. 3c-f). We then visualized single-cell data using UMAP projections of feature counts and observed that cells from the two lines can be perfectly distinguished based on Pol2S5p peaks, H3K27me3 peaks or the combination of the two (Fig. 4a-c). We compared the number of fragments mapping to Pol2S5p and H3K27me3 peaks and observed a strong correlation in both cell types (Fig. 4d, correlation: 0.95), and an even balance of fragments between the two targets in individual cells. In line with this observation, the 400 most variable Pol2S5p peaks and H3K27me3 peaks were sufficient to distinguish the majority of the two cell types (Fig. 4e), demonstrating that CUT&Tag2for1 can be used to identify both active and repressive chromatin features in the same single cells, and we can use them coordinately to distinguish cell identity. To examine the robustness of the single cell CUT&Tag2for1 method as well as the 2for1separator algorithm we performed additional replicates of H1 and K562 cells run on the ICELL8. K562 cells from both replicates cluster nicely together in UMAP space and away from a second cluster composed of H1-hESCs from both replicates according to the deconvolved signal of either Pol2S5p or H3K27me3 (Additional file 4a-b). In addition, we found that cell types from one replicate can be accurately distinguished from one another based on the regions defined as Pol2S5p or H3K27me3 when 2for1separator is applied to an alternative replicate. We conclude that the combination of CUT&Tag2for1 with the 2for1separator algorithm is a robust strategy to identify both active and repressive chromatin features in the same single cells and to coordinately distinguish cell identity.

**Figure 4:**
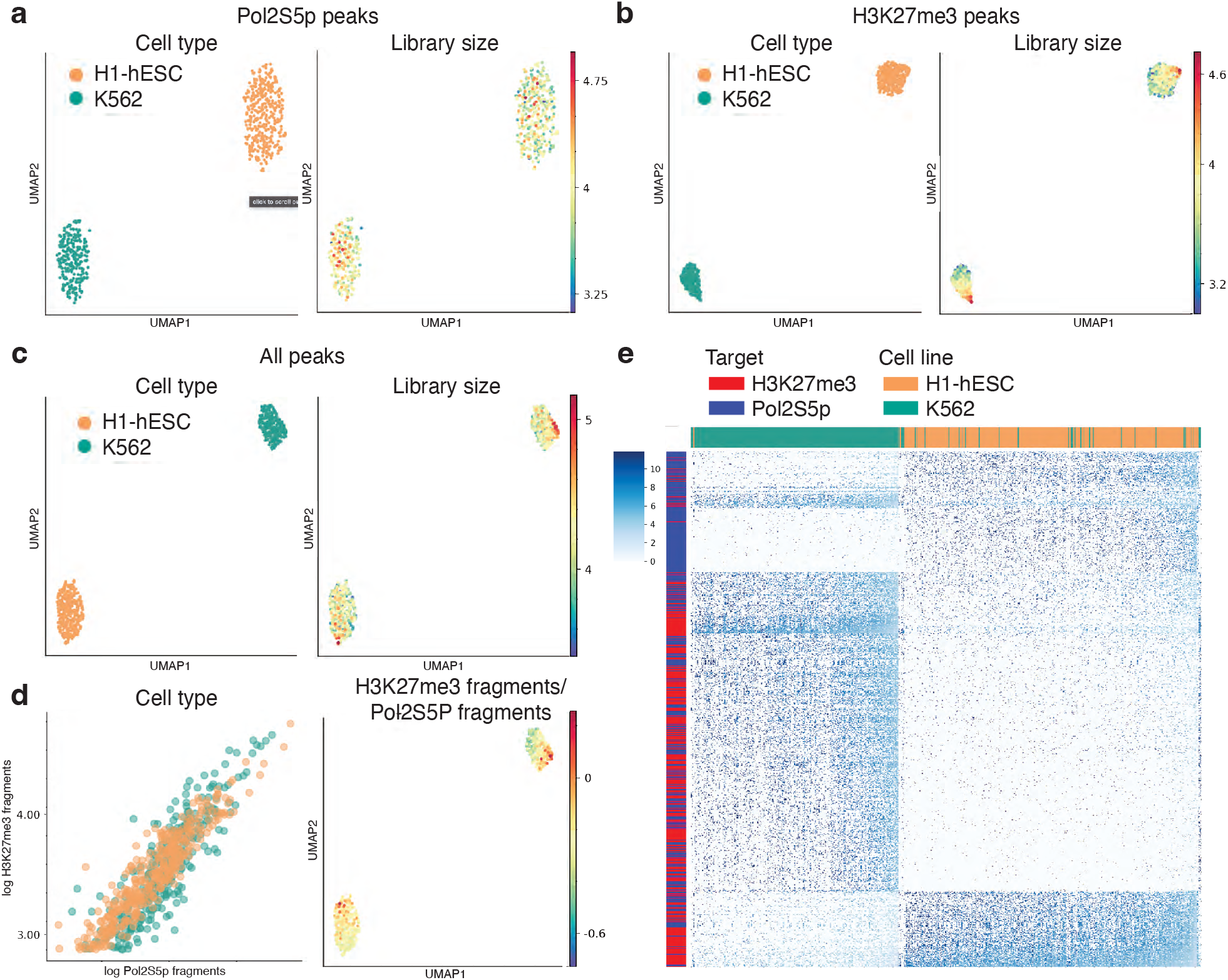
Single-cell CUT&Tag2for1: (**a**) UMAPs representing the low dimensional embedding of cells using inferred Pol2S5p peaks (left), colored by the log of the library size of single cells. (**b**) Same as (a), using inferred H3K27me3 peaks. (**c**) Same as (a), using both inferred Pol2S5p and H3K27me3 peaks. (**d**) Plot comparing the number of fragments mapping to Pol2S5p and H3K27me3 peaks for H1 and K562 cells (left). UMAP of cells colored by fraction of fragments mapping to H3K27me3 peaks (right). (**e**) Heatmap of normalized fragment counts for the top 400 most variable Pol2S5p and H3K27me3 peaks (as rows) from Principal Component 1 across 515 K562 cells and 427 H1 cells (as columns). Results are shown for replicate 1.

## Discussion

Single cell genomics methods for profiling the transcriptome, proteome, methylome and accessible chromatin landscape have advanced rapidly in recent years [19]. Currently, approaches for profiling single epigenome targets on their own or in combination with other modalities (e.g. transcriptome) are the state of the art, but methods for simultaneously profiling the active versus repressive chromatin landscape in single cells are still lacking. CUT&Tag2for1 combines simple antibody mixing in a single workflow with a single sequencing readout to profile and computationally separate accessible and repressed chromatin regions. Single-cell CUT&Tag2for1 avoids the complex workflows, multi-level barcoding and apples-and-oranges integration challenges posed by multimodal profiling methods.

CUT&Tag2for1 was inspired by the observation that Pol2S5p CUTAC, developed based on previously reported H3K4me2 CUTAC [10], yields a different average fragment size profile than H3K27me3 CUT&Tag, and therefore that the two could be distinguished in a single assay. Although the low-salt conditions result in low artifactual accessible site signals in H3K27me3 CUT&Tag experiments [20], in CUT&Tag2for1 these accessible sites accumulate high signals. The DNA fragment length data dimension allows for *a priori* assignment of target origin, which is in keeping with the myriad advantages of using fragment length to elucidate fine grain chromatin structure [21–24]. By also using feature width information in a probabilistic model, we obtain robust separation of the active and repressive landscapes.

Single-cell CUT&Tag2for1 can assign Pol2S5p or H3K27me3 target origin with high fidelity in the absence of ground truth datasets. A limitation of our deconvolution strategy is that it requires a large fragment/large feature and small fragment/small feature pair for best performance. For example, H3K36me3, which marks active gene bodies, might in principle be combined with transcription factors. However, the relatively high abundance of both H3K27me3 and Pol2S5p and the fact that in combination they profile virtually the entire chromatin developmental regulatory landscape, makes our current implementation of CUT&Tag2for1 an attractive genomics-based strategy for a wide range of development and disease studies.

## Acknowledgements

We thank Christine Codomo, Trizia Llagas and Terri Bryson for technical assistance, Jorja Henikoff and Matthew Fitzgibbon for data processing, and the Fred Hutch Shared Genomics and Bioinformatics Resource for DNA sequencing and data handling. This work was supported by grants from the National Institutes of Health to S.H. (R01 HG010492) and to the Fred Hutch Scientific Computing Infrastructure (S10OD028685), and by the Howard Hughes Medical Institute.

## Author Contributions

S.H. conceived CUT&Tag2for1, D.J., K.A. and S.H. performed the experiments, D.O., M.P.M. and M.S. conceived, wrote and implemented the software, M.S. and S.H. provided funding and critical oversight of experiments and analyses, and all authors contributed to writing and editing the manuscript.

## Declaration of Interests

S.H has filed patent applications related to this work.

## CONTACT FOR REAGENT AND RESOURCE SHARING

Further information and requests for resources and reagents should be directed to the Lead Contact, Steven Henikoff

## Methods

### Human Cell culture

Human female K562 Chronic Myleogenous Leukemia cells (ATCC) were authenticated for STR, sterility, human pathogenic virus testing, mycoplasma contamination, and viability at thaw. H1 (WA01) male human embryonic stem cells (hESCs) (WiCell) were authenticated for karyotype, STR, sterility, mycoplasma contamination, and viability at thaw. Cells were cultured as previously described [25]. Briefly, K562 cells were cultured in liquid suspension. H1 cells were cultured in Matrigel (Corning)-coated plates at 37°C and 5% CO_2_ using mTeSR-1 Basal Medium (STEMCELL Technologies) exchanged every 24 hours.

### Bulk CUT&Tag, CUTAC and CUT&Tag2for1

CUTAC using Pol2S5p for accessible site mapping was performed as described in a step-by-step protocol [11]. Briefly, cells were harvested by centrifugation, washed with PBS and nuclei prepared and lightly cross-linked (0.1% formaldehyde 2 min), then washed and resuspended in Wash buffer (10 mM HEPES pH 150 mM NaCl, 2 mM spermidine and Roche complete EDTA-free protease inhibitor), aliquoted with 10% DMSO and slow-frozen to −80°C in Mr. Frosty containers (Sigma-Aldrich cat. no. C1562). CUT&Tag, CUTAC and CUT&Tag2for1 were performed in parallel in single 0.6 mL PCR tubes by mixing with Concanavalin A magnetic beads and performing incubation and wash steps on a magnet. Primary (anti-rabbit) antibody [1:50 for Pol2S5p (Cell Signaling Technology cat. no. 13523) or 1:100 for H3K27me3 (Cell Signaling Technology cat. no. 9733)] in Wash buffer + 0.1% BSA was added and beads were incubated at room temperature for 1-2 hr or overnight at 4 °C. For CUT&Tag2for1, primary antibodies were mixed in the same concentrations. Beads were magnetized and the supernatant was removed, then the beads were resuspended in guinea pig anti-rabbit secondary antibody (Antibodies Online cat. no. ABIN101961) and incubated 0.5-1 hr. Beads were magnetized, the supernatant was removed, then the beads were resuspended in pAG-Tn5 pre-loaded with mosaic-end adapters (Epicypher cat. no. 15-1117 1:20) in 300-wash buffer (Wash buffer except containing 300 mM NaCl) and incubated 1-2 hr at room temperature. Following magnetization, supernatant removal and washing in 300-wash buffer, the beads were incubated at 37 °C in either 10 mM MgCl_2_, 300 mM NaCl (for CUT&Tag) for 1 hr or 5 mM MgCl_2_, 10 mM TAPS pH 8.5 (for CUTAC and CUT&Tag2for1) for 10-30 min. In some experiments CUTAC and CUT&Tag2for1 tagmentation was performed in 5 mM MgCl_2_, 10 mM TAPS pH 8.5 with addition of 10% (w/v) 1,6-hexanediol (Sigma-Aldrich cat. no. 240117-50G) or 10% (v/v) N,N-dimethylformamide [10]. Bead suspensions were chilled on ice, magnetized, the supernatant was removed, then beads were washed with 10 mM TAPS pH 8.5, 0.2 mM EDTA, and resuspended in 5 µL 0.1% SDS, 10 mM TAPS pH 8.5. Beads were incubated at 58 °C in a thermocycler with heated lid for 1 hr, followed by addition of 15 µL 0.67% Triton X-100 to neutralize the SDS. Barcoded PCR primers were added followed by 25 µL of either NEBNext 2x Master Mix (ME541L, non-hotstart) or KAPA Polymerase 2x master mix [Roche KAPA HiFi plus dNTPs: 360 µL 5X KAPA HiFi buffer, 54 µL KAPA dNTP mix (10 mM each), 36 µL KAPA non-hotstart DNA Pol (1U/µL), 450 µL dH_2_O]. Gap-filling and 12-cycle PCR were performed: 58 °C 5 min, 72 °C 5 min, 98 °C 30 sec, 12 cycles of (98 °C 10 sec denaturation and 60 °C 10 sec annealing/extension), 72 °C 1 min, and 8 °C hold. In some experiments linear pre-amplification was performed using this program with 3-12 cycles but with only i5 primers, followed by addition of i7 primers at 8 °C and 10-12 cycles of (98 °C 10 sec denaturation and 60 °C 10 sec annealing/extension), then 72 °C 1 min, and 8 °C hold, and in other experiments the initial 98 °C denaturation step was extended from 30 sec to 5 min, but no consistent differences in the resulting libraries were observed. SPRI paramagnetic beads were added directly to the bead-cell slurry for clean-up as described by the manufacturer (Magbio Genomics, cat. no. AC-60500). Elution was in 20 µL 1 mM Tris pH 8.0, 0.1 mM EDTA. Library quality and concentration were evaluated by Agilent Tapestation capillary gel analysis, barcoded libraries were mixed and PE25 sequencing performed on an Illumina HiSeq2500 by the Fred Hutch Genomics Shared Resource.

### Single-cell CUT&Tag2for1

CUT&Tag2for1 was performed using lightly fixed K562 and H1 nuclei. Frozen nuclei were thawed and aliquots containing 20,000 nuclei were centrifuged at 700 x g for 4 minutes at 4 °C. Nuclei were washed once with Wash buffer, centrifuged again, and then resuspended in Antibody buffer (10 mM HEPES pH 150 mM NaCl, 2 mM spermidine, 2 mM EDTA, 0.1% BSA, and Roche complete EDTA-free protease inhibitor) with primary anti-Pol2S5p antibody (Cell Signaling Technology cat. no. 13523, 1:50) and anti-H3K27me3 (Cell Signaling Technology cat. no. 9733, 1:100) in 0.6 mL PCR tubes. Primary antibody binding was performed overnight at 4 °C. Samples were centrifuged at 700 x g for 4 minutes at 4 °C between incubation steps. and incubated for 1 hour at room temperature for the guinea pig anti-rabbit secondary antibody (Antibodies Online cat. no. ABIN101961 1:100) and for 1 hour at room temperature for pAG-Tn5 (Epicypher cat. no. 15-1117, 1:20) tethering. Samples were then centrifuged, washed with 300-wash buffer, pelleted by centrifugation, and then resuspended in 5 mM MgCl_2_, 10% hexanediol, 10 mM TAPS pH 8.5 for 20 min at 37 °C for tagmentaion. Reactions were stopped by adding EDTA to a final concentration of 1 mM, and kept at 4 °C until dispensation on the ICELL8 platform.

Cells were processed on the ICELL8 instrument according to a previously optimized protocol for release of tagmented DNA in SDS, followed by a Triton X-100 neutralization step and PCR amplification [26]. Briefly, the volume of 10 mM TAPS Buffer pH 8.5 was adjusted to 65 µL per 20,000 nuclei to yield a concentration of ~300 nuclei/µL and nuclei were stained with 1X DAPI and 1X secondary diluent reagent (Takara Cat# 640196). The 8 source wells of the ICELL8 were loaded with 65 µL of the suspension of tagmented nuclei and dispensed into a SMARTer ICELL8 350v chip (Takara Bio, cat. no. 640019) at 35 nL per well. The chip was then sealed for imaging and spun down at 1200 x g for 1 min. Imaging on a DAPI channel confirmed the presence of single cells in specific wells. Non-single-cell wells were excluded from downstream reagent dispenses. A volume of 35 nL of 0.19% SDS in 10 mM TAPS Buffer pH 8.5 was dispensed into active wells and the chip was dried, sealed and spun down at 1200 x g for 1 min. The chip was placed in a thermocycler and held at 58 °C for 1 hr to release tagmented chromatin. The chip was spun at 1200 x g for 1 min before opening, and 35 nL of 2.5% Triton X-100 in distilled deionized H_2_0 was dispensed into all active wells. To index the whole chip, 72 × 72 i5/i7 primers containing unique indices (5,184 microwells total) were dispensed at 35 nL in wells that contained single cells, followed by two dispenses of 50 nL (100 nL total) KAPA PCR mix (2.775 X HiFi Buffer, 0.85 mM dNTPs, 0.05 U KAPA HiFi polymerase / µL, Roche Cat# 07958846001). The chip was sealed for heated incubation and spun down at 1200 x g for 1 min after each dispense. PCR on the chip was performed with the following protocol: 5 min at 58 °C, 10 min at 72 °C and 2 min at 98°C, followed by 15 cycles of 15s at 98 °C, 15s at 60 °C and 10s at 72 °C, with a final extension at 72 °C for 2 min. The contents of the chip were then centrifuged into a collection tube (Takara Cat# 640048) at 1200g for 3 min. Two rounds of SPRI bead cleanup at a 1.3 : 1 v/v ratio of beads to sample were performed to remove residual PCR primers and detergent. Samples were resuspended in 20 µL of 10 mM Tris-HCl pH 8.0. Library quality and concentration were evaluated by Agilent Tapestation capillary gel analysis, and single-cell CUT&Tag2for1 samples were then pooled with bulk libraries prepared using compatible barcodes. PE25 sequencing was performed on an Illumina HiSeq2500 by the Fred Hutch Genomics Shared Resource.

### Deconvolution using fragment size (bulk)

Peaks were called using SEACR v1.3 [27]. Fragments overlapping peaks were ascertained using bedtools intersect [28]. For each peak, we calculated the average fragment size of all fragments overlapping the peak in question, and then fit the distribution of average fragment sizes across all peaks to a mixture of two Gaussian distributions using Mixtools NormalMixEM [18]. Peaks were partitioned into “large” (H3K27me3) and “small” (PolIIS5P) fragment size classes based on the average fragment size threshold at which the two calculated Gaussian distributions intersect. Bulk H3K27me3 CUT&Tag and PolIIS5P CUTAC were mapped onto large and small peak classes in heatmap form using Deeptools [29].

### Deconvolution using feature width and fragment size (single-cell)

CUT&Tag fragments result from two independent integration events resulting in two tagmentation cut sites after gap-filling, barcoded PCR and DNA sequencing. Rather than trying to attribute each fragment to either Pol2S5p or H3K27me3, our deconvolution approach estimates how likely was a cut derived from Pol2S5p or H3K27me3 antibodies. We use three key insights for deconvolution: (i) fragment length distributions are significantly different between the two targets (Additional file 1), (ii) cuts from a target have a positional dependency *i*.*e*., observation of multiple cuts from a specific target at a genomic location most likely means a cut close to this set was induced by the same target, and (iii) feature widths between the two targets are typically different. Pol2S5p peaks are narrow and sharp, whereas H3K27me3 domains are broad and diffuse (Fig. 2b). Motivated by this, we developed 2for1separator, an algorithm to deconvolve the CUT&Tag2for1 data into two signal tracks – representing the density of chromatin cut sites targeted by H3K27me3 and PolS5p antibodies respectively. We then use the deconvolved signals to identify narrow Pol2S5p peaks and broad H3K27me3 domains in the data.

### Overview of 2for1separator

Formally, we represent a cut as a tuple (*x, l*) where *x* stands for the location in the genome and *l* the length of the fragment it belongs to. The density of CUT&Tag2for1 cuts at cut-site *x* with fragment length, *l* can be represented as

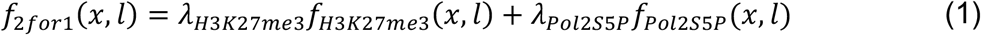

where function *f* is the probability density function (PDF). *λ*_*H*3*K*27*me*3_ and *λ*_*Pol*2*S*5*P*_ represent the respective weights.

We assume that the length *l* and position *x* are independently distributed for each target, therefore *f*_*H*3*K*27*me*3_(*x, l*) can be written as

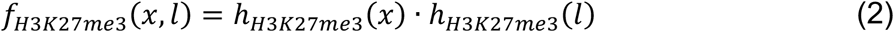

Similarly, for Pol2S5p

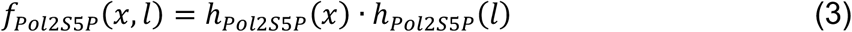

where *h*(*x*) is the location-specific marginal cut-side probability density function and *h*(*l*) is the location-independent marginal fragment length probability density function.

### Fragment length distribution prior

The fragment length marginal PDFs, *h*_*Pol*2*S*5*P*_(*l*) and *h*_*H*3*K*2*7me*3_(*l*), are parameterized separately to account for the differences in length distributions between the two targets. Length distributions show characteristic modes irrespective of the target (Additional files 1b-c, 2a-b). We thus represent the fragment length PDF as a mixture of four log-normal distributions with modes centered at 70, 200, 400, 600 (Fig. 3b). We do not make a distinction for fragments that are >800 base pairs in length since they occur rarely. We assume the weights of the modes to follow a Dirichlet distribution – effective for modeling multinomial distributions, that we roughly based on the single antibody data.

Through a rough estimate of these mode weights and with arguable uncertainty of the true distribution we came to use the Dirichlet-parameter vector (450, 100, 10, 1) for Pol2S5p and (150, 300, 50, 10) for H3K27me3. Note that the deconvolution inferred weights remain very consistent across multiple fragment subsamples while deviating strongly from the mean of the Dirichlet prior (Additional file 2c-d), indicating that the result is data driven and not very sensitive to the exact choice of prior parameters. We only need to encode the fact that Pol2S5p fragments are shorter on average than H3K27me3 fragments.

### Cut-site densities and prior

We chose to model the cut-site PDFs as Gaussian Processes (GP), a powerful technique that can accurately infer the shape of the signal by considering the positional dependencies in signal values (Fig. 3b). The GP is used to predict the log cut density at a particular cut-site as a function of all the cuts in the neighborhood. A GP is defined by mean and covariance functions where the covariance function encodes the neighborhood information, *i*.*e*., positional dependencies between cuts and feature widths, making GPs ideally suited to infer cut-site density functions for the two targets.

We took an empirical approach to define the covariance function of the Gaussian process. We examined the Gaussian kernel density estimates (***σ*** = 200) of cuts from the H3K27me3 CUT&Tag and Pol2S5p CUTAC experiments and determined that the autocorrelation of the log-density, representing both local dependencies, is well approximated through the Matérn covariance function (*nu* = 3/2) [30]. Based on the observed autocorrelations, we chose this covariance function with length scales 500 and 2000 as kernels of the GP for the Pol2S5p and H3K27me3 targets respectively to account for feature width differences. We also note that differences in feature widths is not a necessary component, and our model can deconvolve the signals as long as the fragment length distributions between the two targets are different.

### Constraints on the Gaussian Process

The functions generated through the GP express the desired smoothness and mean value but are not guaranteed to represent probability density functions. To ensure that the generated functions indeed represent PDFs, we must guarantee two additional constraints: (i) strict positivity (ii) a fixed integral, without which, the resulting likelihood could grow infinitely jeopardizing any posterior estimate of the location specific PDFs.

Positivity is ensured by applying the exponential: We model the cut-site PDF *h*_*POl2S5P*_(*x*) as

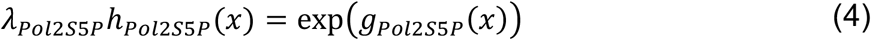

where *g*_*Pol*2*S*5*P*_ is a random variable of a GP. Similarly,

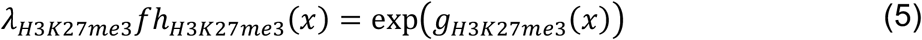

The sum of the two PDFs in Equations (4) and (5) should integrate to one for a fixed integral. Rather than constraining the integral to one, we aim for a density function that integrates to the total number of observed cuts for ease of implementation. This representation results in a constant factor in the combined likelihood function and does not impact the inference. As an added benefit of this formulation, the inferred density function has the unit “cuts per base pair” and hence is insensitive to the size of the deconvolved genomic region. Further, this also results in the log-density to have an approximate mean value of 0 across the whole genome and thus we use a zero-mean GP. We approximate this integral with the rectangle rule, by assuming one rectangle per cut site and a width such that neighboring rectangles touch at the midpoint between the cut sites. To enforce the correct integral, we impose a log-normal distribution of the resulting approximation around the desired value and a very small standard deviation of 0.001, since enforcing a constraint to a fixed value makes the inference intractable.

### Inference

To infer the most likely target specific chromatin cut PDF, we use the gradient descent method, limited-memory BFGS on the posterior parameter distribution to find the local maximum a posteriori point (MAP). The MAP represents the most likely cut PDFs and fragment length distributions in the chosen parametrization of our model.

### Pol2S5p peak calling

We use the deconvolved Pol2S5p signals to perform peak calling. We nominate each region containing cuts with deconvolved Pol2S5p signal greater than a computed threshold as Pol2S5p peaks. We retain Pol2S5P peaks longer than 100 bases for downstream analysis. We identify the position within the peak with maximal deconvolved signal as the summit.

We first estimate the fraction of cuts that are derived from Pol2S5p to compute the threshold. The fraction, denoted as 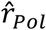, is estimated as the ratio between the integral of Pol2S5p deconvolved density and the integral of the combined density. In practice, we found this estimate to be susceptible to instability and we therefore used a beta distribution with parameters *α* = 0.5, *β* = 0.5 as a prior to derive a robust estimate. With a further conservative assumption that 50% of the Pol2S5p cuts fall into Pol2S5p reproducible peaks, the expected value of the fraction of cuts that fall in Pol2S5p peaks is

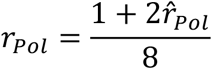

We therefore use the *r*_*Pol*_th percentile of the deconvolved signal value as the threshold i.e., regions with cuts with deconvolved signal higher than the *r*_*Pol*_th percentile are identified as Pol2S5p peaks.

### H3K27me3 domains

A procedure analogous to Pol2S5p peak calling was used to identify H3K27me3 domains using the deconvolved H3K27me3 signal. We observed that large H3K27me3 domains appear as discontinuous signal blocks (Fig. 3c – right panel). We therefore applied an additional smoothing on the deconvolved H3K27me3 signal using a Gaussian filter and computed the average between the smoothed and original signal. We then repeated the peak calling procedure on the smoothed signal and identified H3K27me3 domains as the union of domains identified using deconvolved and the additionally smoothed signals. Only peaks wider than 400 bases are retained for downstream analysis.

### Overlap peaks

A fraction of genomic sites were identified as peaks in both Pol2S5p and H3K27me3. If the overlaps of a H3K27me3 peak with Pol2S5p is less than 50% of the H3K27me3 peak span, we resolve the region as a H3K27me3 peak (and vice-versa for Pol2S5p peaks). The remainder of the peaks are termed as overlap peaks (Additional file 5). Overlap peaks comprise only ~1% of the total (139 of 10,661 H1 peaks and 104 of 11,111 K562 peaks) and were not used in the analysis.

### Implementation Details

Since the GP employs the covariance between all cut-sites, the memory demand grows approximately quadratic with the number of unique cut-sites. However, cuts that are further apart than 10,000 base pairs express no relevant covariance and must not be considered in the same GP. We use this observation to split up genomic regions into intervals with at most 10,000 unique cut sites. We pad each interval with an additional 10,000 bases on either side to ensure stable estimation of the signal at the interval boundaries and discard the padding after deconvolution. A GP is fit separately for each interval and the results concatenated to obtain a deconvolution of all genomic regions.

We also limit the deconvolution to regions where the Gaussian kernel density estimate of all cuts (bandwidth=200) indicates at least 2 cuts per 100 base pairs. Neighboring regions are merged if they are separated by fewer than 10,000 base pairs. These selected regions were segmented into intervals as described above. We then grouped all intervals of the selected regions into ~200 tasks and applied the posteriori point maximization of PyMC3 [31] for deconvolution.

### Application to single-cell data

We aggregated the reads of all cells of the two cell types from both the replicates into a pseudo-bulk set of fragments for each cell type. After applying our 2for1separator algorithm to identify Pol2S5p and H3K27me3 peaks we used featureCount [32] to count the number of fragments that overlap each peak for each cell and target. We analyzed the two replicates separately to avoid misleading batch effects.

We binarized the matrix and normalized the data using TF-IDF following the ArchR solution [33], separately for H3K27me3 and Pol2S5p. We then applied Singular Value Decomposition to the normalized and log-transformed data and used 30 components for downstream analysis. Since the first or second principal component remains very strongly correlated with the library size despite the normalization, we exclude the respective component in the UMAP and other downstream analyses.

Batch correction between the two replicates was performed using Harmony [34]. For independent replicate analysis in Additional file 3c-d, deconvolution was performed separately for each replicate and peak counts from one were used to compute visualizations and clustering for the other replicate.

## Data and code availability

The 2for1separator algorithm for single-cell CUT&Tag2for1 deconvolution is available at https://github.com/settylab/2for1separator.

Other software is provided at https://github.com/mpmeers/MeersEtAl_MulTI-Tag and https://github.com/FredHutch/SEACR.

## Notes

### Competing Interest Statement

S.H. has filed patent applications on related work.

## References

1. Waddington C: How animals develop. George Allen & Unwin; 1946.

2. Simon JA, Kingston RE: Occupying chromatin: Polycomb mechanisms for getting to genomic targets, stopping transcriptional traffic, and staying put. Mol Cell 2013, 49:808–824.

3. Klein DC, Hainer SJ: Genomic methods in profiling DNA accessibility and factor localization. Chromosome Res 2020, 28:69–85.

4. Kaya-Okur HS, Wu SJ, Codomo CA, Pledger ES, Bryson TD, Henikoff JG, Ahmad K, Henikoff S: CUT&Tag for efficient epigenomic profiling of small samples and single cells. Nature communications 2019, 10:1930.

5. Wu SJ, Furlan SN, Mihalas AB, Kaya-Okur HS, Feroze AH, Emerson SN, Zheng Y, Carson K, Cimino PJ, Keene CD, et al: Single-cell CUT&Tag analysis of chromatin modifications in differentiation and tumor progression. Nat Biotechnol 2021, 39:819–824.

6. Bartosovic M, Kabbe M, Castelo-Branco G: Single-cell CUT&Tag profiles histone modifications and transcription factors in complex tissues. Nat Biotechnol 2021, 39:825–835.

7. Zhu C, Zhang Y, Li YE, Lucero J, Behrens MM, Ren B: Joint profiling of histone modifications and transcriptome in single cells from mouse brain. Nat Methods 2021, 18:283–292.

8. Regev A, Teichmann SA, Lander ES, Amit I, Benoist C, Birney E, Bodenmiller B, Campbell P, Carninci P, Clatworthy M, et al: The Human Cell Atlas. Elife 2017, 6:e27041.

9. Zhu C, Yu M, Huang H, Juric I, Abnousi A, Hu R, Lucero J, Behrens MM, Hu M, Ren B: An ultra high-throughput method for single-cell joint analysis of open chromatin and transcriptome. Nat Struct Mol Biol 2019, 26:1063–1070.

10. Henikoff S, Henikoff JG, Kaya-Okur HS, Ahmad K: Efficient chromatin accessibility mapping in situ by nucleosome-tethered tagmentation. Elife 2020, 9:e63274.

11. Henikoff S, Henikoff JG, Ahmad K: Simplified Epigenome Profiling Using Antibody-tethered Tagmentation. bio-protocol 2021, 11:e4043.

12. Landt SG, Marinov GK, Kundaje A, Kheradpour P, Pauli F, Batzoglou S, Bernstein BE, Bickel P, Brown JB, Cayting P, et al: ChIP-seq guidelines and practices of the ENCODE and modENCODE consortia. Genome Res 2012, 22:1813–1831.

13. Buenrostro JD, Giresi PG, Zaba LC, Chang HY, Greenleaf WJ: Transposition of native chromatin for fast and sensitive epigenomic profiling of open chromatin, DNA-binding proteins and nucleosome position. Nat Methods 2013, 10:1213–1218.

14. Mahat DB, Kwak H, Booth GT, Jonkers IH, Danko CG, Patel RK, Waters CT, Munson K, Core LJ, Lis JT: Base-pair-resolution genome-wide mapping of active RNA polymerases using precision nuclear run-on (PRO-seq). Nat Protoc 2016, 11:1455–1476.

15. Soares LM, He PC, Chun Y, Suh H, Kim T, Buratowski S: Determinants of Histone H3K4 Methylation Patterns. Mol Cell 2017, 68:773–785 e776.

16. Andersson R, Sandelin A, Danko CG: A unified architecture of transcriptional regulatory elements. Trends Genet 2015, 31:426–433.

17. Meshorer E, Yellajoshula D, George E, Scambler PJ, Brown DT, Misteli T: Hyperdynamic plasticity of chromatin proteins in pluripotent embryonic stem cells. Dev Cell 2006, 10:105–116.

18. Benaglia T, Chauveneau D, Hunter DR, Young DS: mixtools: An R Package for Analyzing Mixture Models. J Statistical Software 2010, 32.

19. Zhu C, Preissl S, Ren B: Single-cell multimodal omics: the power of many. Nat Methods 2020, 17:11–14.

20. Kaya-Okur HS, Janssens DH, Henikoff JG, Ahmad K, Henikoff S: Efficient low-cost chromatin profiling with CUT&Tag. Nat Protoc 2020, 15:3264–3283.

21. Kent NA, Adams S, Moorhouse A, Paszkiewicz K: Chromatin particle spectrum analysis: a method for comparative chromatin structure analysis using paired-end mode next-generation DNA sequencing. Nucleic Acids Res 2011, 39:e26.

22. Henikoff JG, Belsky JA, Krassovsky K, Macalpine DM, Henikoff S: Epigenome characterization at single base-pair resolution. Proc Natl Acad Sci U S A 2011, 108:18318–18323.

23. Meers MP, Janssens DH, Henikoff S: Pioneer Factor-Nucleosome Binding Events during Differentiation Are Motif Encoded. Mol Cell 2019, 75:562–575.

24. Ramachandran S, Henikoff S: Transcriptional Regulators Compete with Nucleosomes Post-replication. Cell 2016, 165:580–592.

25. Janssens DH, Wu SJ, Sarthy JF, Meers MP, Myers CH, Olson JM, Ahmad K, Henikoff S: Automated in situ chromatin profiling efficiently resolves cell types and gene regulatory programs. Epigenetics Chromatin 2018, 11:74.

26. Janssens DH, Meers MP, Wu Sj, Babaeva E, Meshinchi S, Sarthy JF, Ahmad K, Henikoff S: Automated CUT&Tag profiling of chromatin heterogeneity in mixed-lineage leukemia. Nat Genet 2021, https://www.nature.com/articles/s41588-021-00941-9

27. Meers MP, Tenenbaum D, Henikoff S: Peak calling by Sparse Enrichment Analysis for CUT&RUN chromatin profiling. Epigenetics Chromatin 2019, 12:42.

28. Quinlan AR, Hall IM: BEDTools: a flexible suite of utilities for comparing genomic features. Bioinformatics 2010, 26:841–842.

29. Ramirez F, Ryan DP, Gruning B, Bhardwaj V, Kilpert F, Richter AS, Heyne S, Dundar F, Manke T: deepTools2: a next generation web server for deep-sequencing data analysis. Nucleic Acids Res 2016, 44:W160–165.

30. Genton MG: Classes of kernels for machine learning: a statistics perspective. J Mach Learn Res 2002, 2:299–312.

31. Salvatier J, Wiecki TV, Fonnesbeck C: Probabilistic programming in Python using PyMC3. PeerJ Computer Science 2016, 2:e55.

32. Liao Y, Smyth GK, Shi W: featureCounts: an efficient general purpose program for assigning sequence reads to genomic features. Bioinformatics 2013, 30:923–930.

33. Granja JM, Corces MR, Pierce SE, Bagdatli ST, Choudhry H, Chang HY, Greenleaf WJ: ArchR is a scalable software package for integrative single-cell chromatin accessibility analysis. Nat Genet 2021, 53:403–411.

34. Korsunsky I, Millard N, Fan J, Slowikowski K, Zhang F, Wei K, Baglaenko Y, Brenner M, Loh PR, Raychaudhuri S: Fast, sensitive and accurate integration of single-cell data with Harmony. Nat Methods 2019, 16:1289–1296.

