## Additional file 2 for "Simultaneous CUT&Tag profiling of the accessible and silenced regulome in single cells"

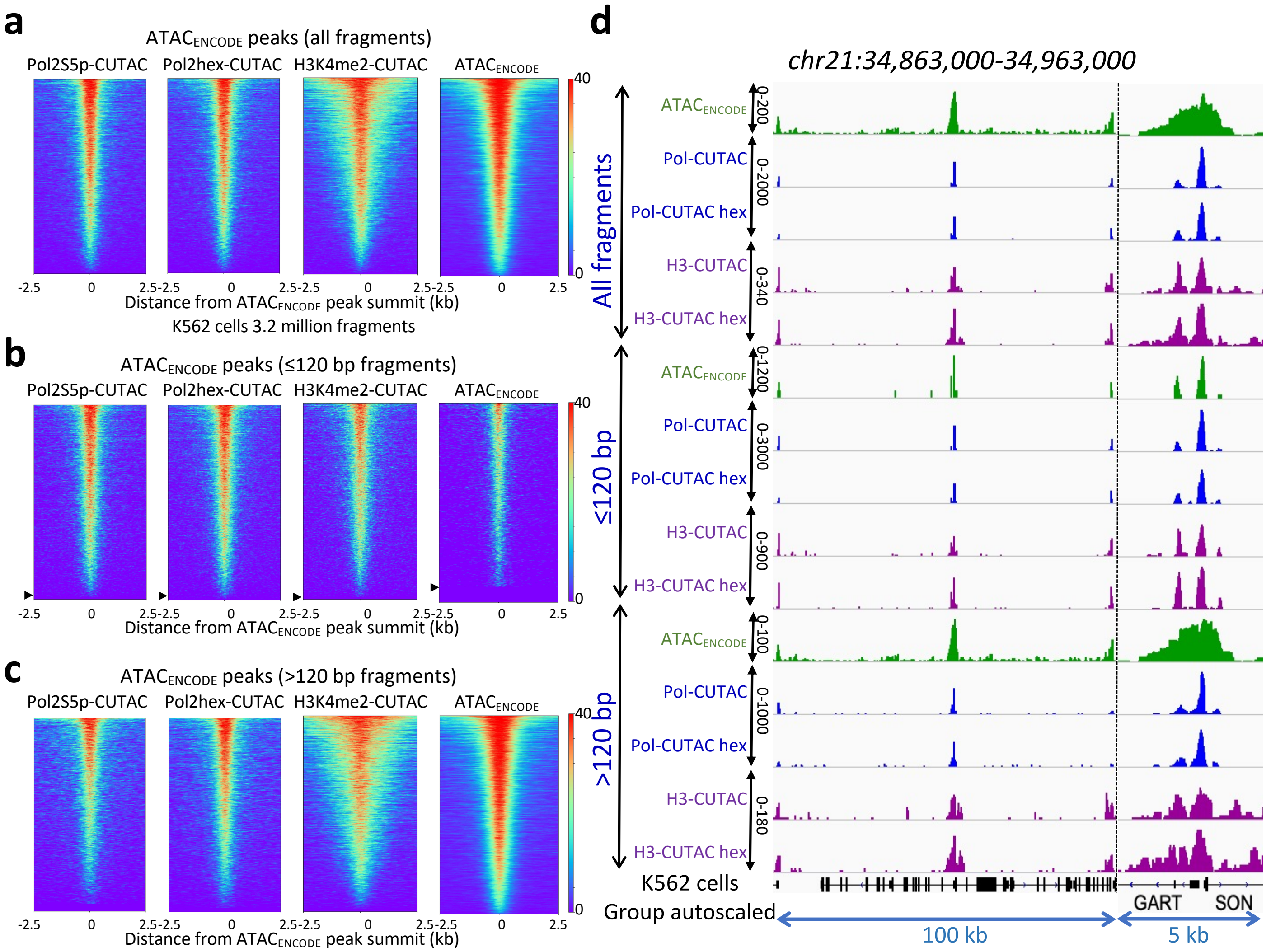


**e**

| N=38155 | ATAC_1 | ATAC_2 | H3-CUTAC | H3-CUTAC hex | Pol-CUTAC | Pol-CUTAC hex |
|---|---|---|---|---|---|---|
| ATAC_1 | 1 | 0.89 | 0.62 | 0.62 | 0.52 | 0.54 |
| ATAC_2 | 0.89 | 1 | 0.61 | 0.62 | 0.52 | 0.53 |
| H3-CUTAC | 0.62 | 0.61 | 1 | 0.9 | 0.66 | 0.65 |
| H3-CUTAC hex | 0.62 | 0.62 | 0.9 | 1 | 0.58 | 0.58 |
| Pol-CUTAC | 0.52 | 0.52 | 0.66 | 0.58 | 1 | 0.96 |
| Pol-CUTAC hex | 0.54 | 0.53 | 0.65 | 0.58 | 0.96 | 1 |

R: 0, 0.5, 1

**Additional file 2: Close correspondences between CUTAC and ATAC-seq** (**a**) Heatmaps showing alignments of Pol2S5p-CUTAC ±10% 1,6-hexanediol during tagmentation, H3K4me2-CUTAC and ATAC$_{ENCODE}$ data over peaks called on an ATAC$_{ENCODE}$ replicated dataset. (**b**) Same as (a) showing the ≤120-bp fragment subset, where arrowheads indicate the row where enrichment of signal over peaks ends: 98-99% for CUTAC and 93% for ATAC$_{ENCODE}$. (**c**) Same as (b) showing the >120 bp subset. Fragments were mapped to hg19, and 3.2 million fragments were randomly sampled from each dataset and used to make bedgraph tracks. **(d**) Tracks comparing Pol2S5p and H3K4me2 CUTAC to ATAC-seq using K562 cell data generated by the ENCODE project. The region around the GART-SON bidirectional promoter is shown at two different scales. (**e**) Correlation matrix of CUTAC and ATAC-seq datasets used in this analysis. ATAC_1 is the ENCODE dataset used to call peaks and ATAC_2 is a replicate of ATAC_1.
