## Additional file 1 for "Simultaneous CUT&Tag profiling of the accessible and silenced regulome in single cells"

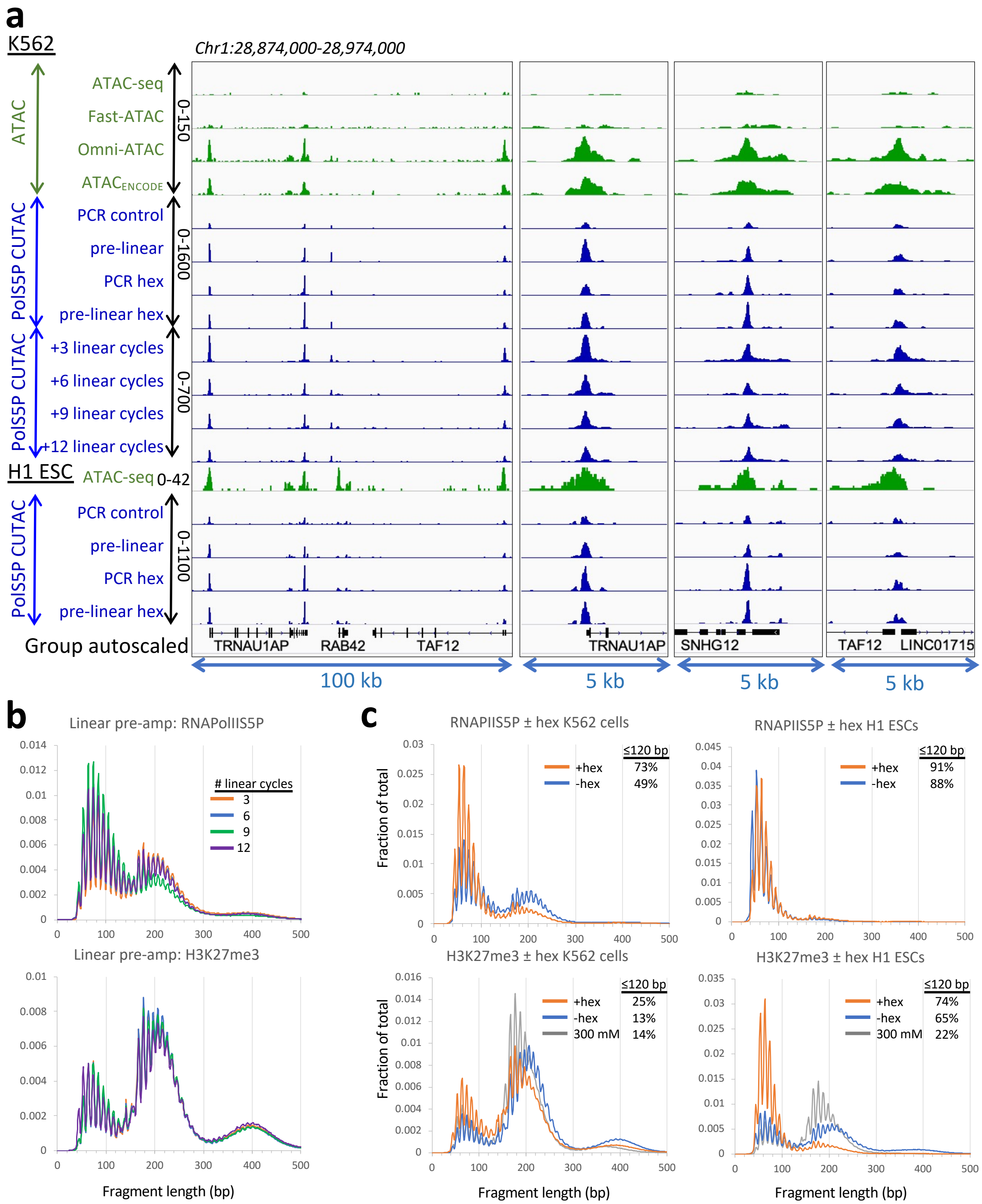


**Additional file 1: Feature definition and fragment length separation under CUTAC conditions. (a)** Fragments were mapped to hg19, and 3.2 million fragments were randomly sampled from each dataset and used to make bedgraph tracks. A representative region is shown. To compare peaks with very different signal-to-noise levels, samples were group-autoscaled with ranges indicated to the left of each set of tracks. Pol2S5p CUTAC of K562 cells with linear pre-amplification using only P5 primers for 12 cycles was followed by addition of P7 primers and PCR for various numbers of cycles. (**b**) Size distributions were not affected by differences in the number of PCR cycles following linear amplification. (**c**) Fragment length distributions for K562 cells (left) and H1 ES cells (right) are shown for linear pre-amplified fragments after tagmentations in 10 mM TAPS + 5 mM $MgCl_2$ ± 1,6-hexanediol or 300 mM NaCl + 10 mM $MgCl_2$ plotted as fractions of the total mapped fragments for linear pre-amplified datasets. Percentages of total fragments ≤120 bp iare shown. Tagmentation in 1,6-hexanediol generally results in a smaller fragment distribution, especially conspicuous for H1 cell nucleosomes. The higher recovery of nucleosome-sized fragments from K562 cells than H1 ES cells reflects the much lower abundance of Polycomb domains in H1 cells.
