## Additional file 3 for "Simultaneous CUT&Tag profiling of the accessible and silenced regulome in single cells"

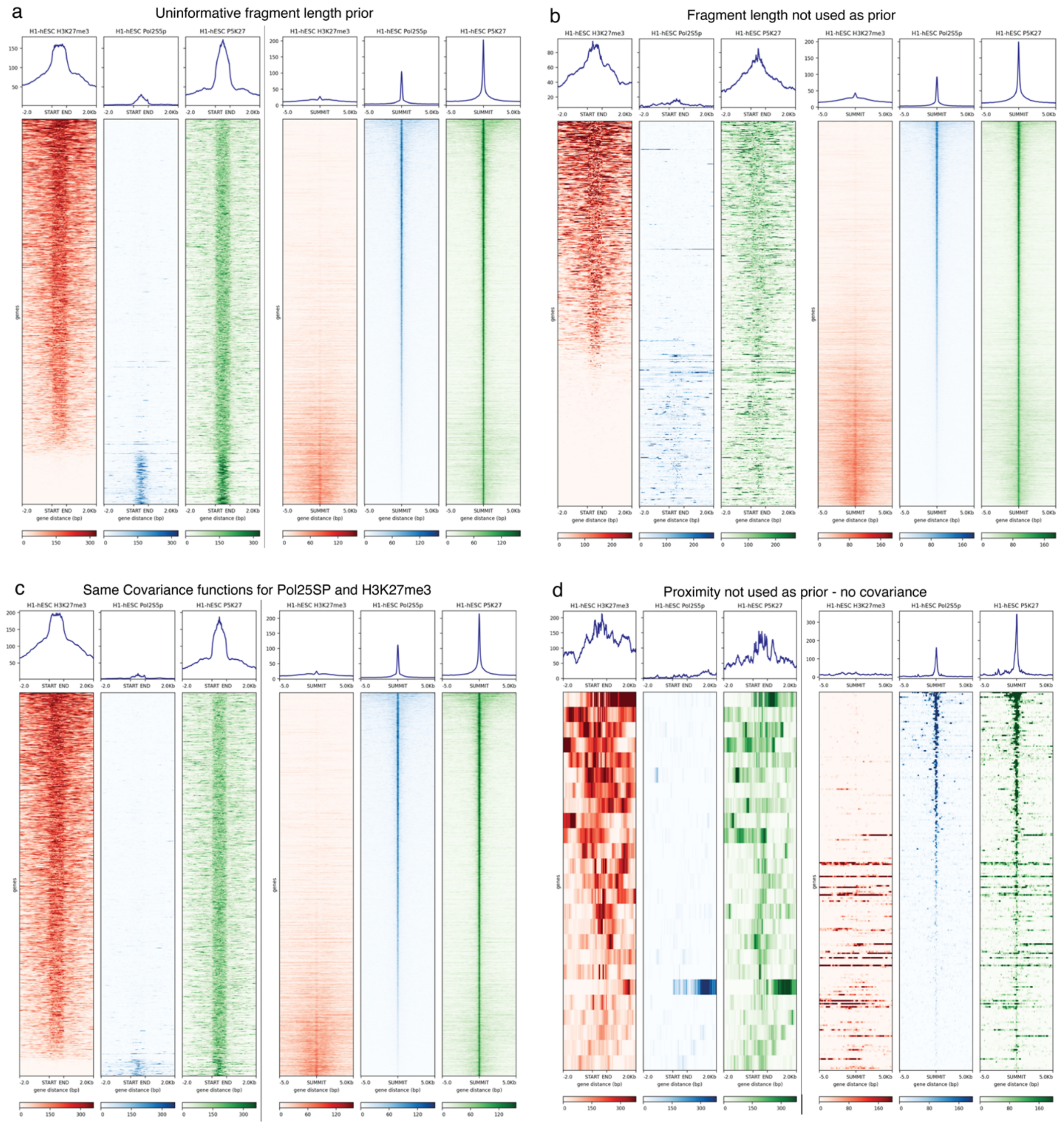


**Additional File 3: Determination of critical features for deconvolution by 2for1separator.** (**a**) Deconvolution of H1 CUT&Tag2for1 data using an uninformative dirichlet prior (1, 1, 1, 1) for fragment length distribution. Left heatmaps show signal for H3K27me3 and right heatmaps show signals for Pol2S5P. Deconvolution is not dependent on prior knowledge about fragment size distributions. (**b**) Deconvolution without fragment length as prior leads to drop in performance. (**c**) Use of same covariance function for H3K27me3 and Pol2S5P does not degrade performance whereas not using proximity information leads to failure of denconvolution (**d**).
