## Additional file 4 for "Simultaneous CUT&Tag profiling of the accessible and silenced regulome in single cells"

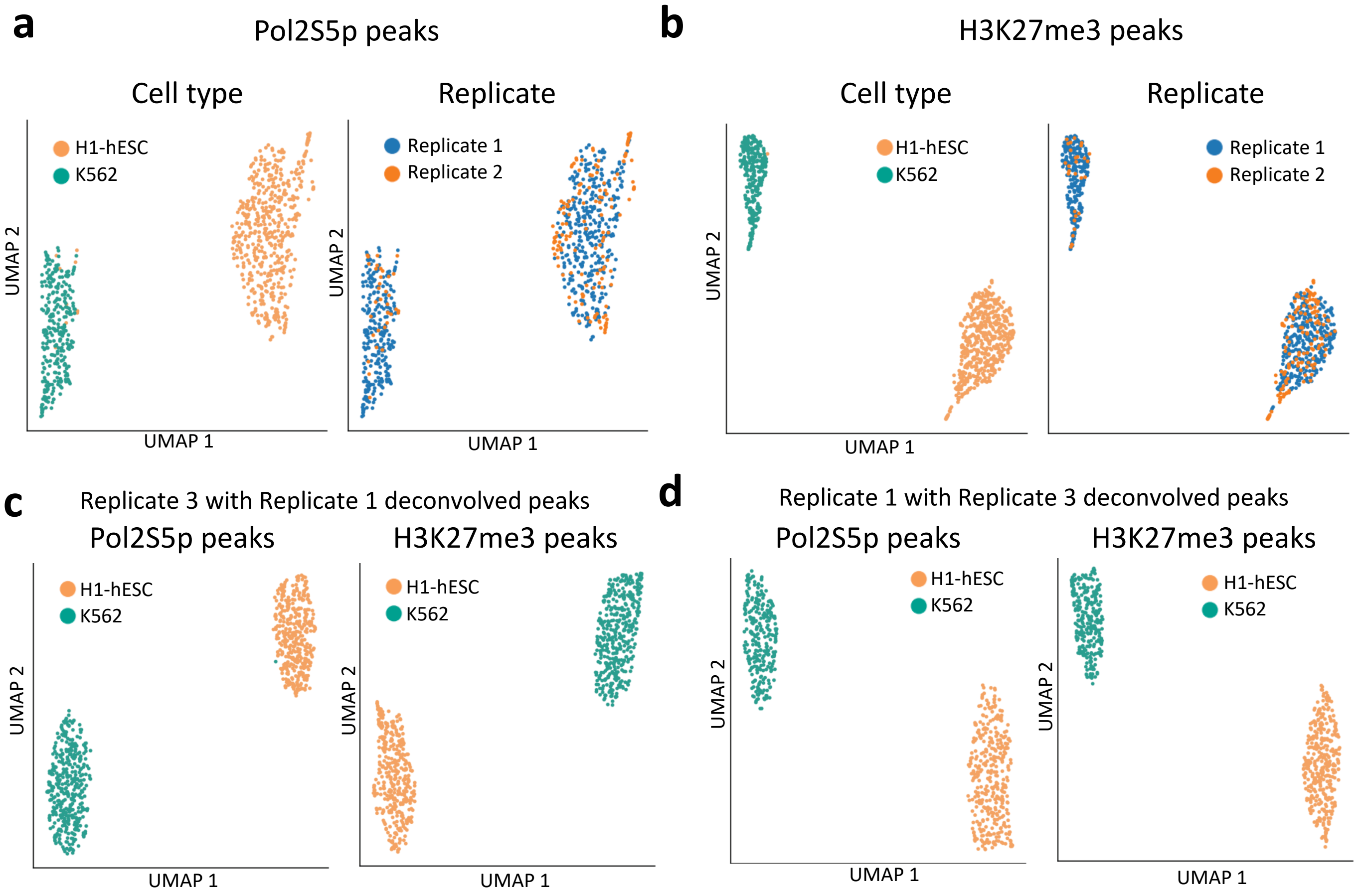


**Additional file 4: Single Cell CUT&Tag2for1 and 2for1separator are consistent across replicates** (**a**) UMAP plot showing that single K562 cells run as independent replicates cluster together according to their deconvolved Pol2S5p signal and away from the two replicates of H1-hESCs. (**b**) Same as (a) showing cells from two independent replicates clustered according to their deconvolved H3K27me3 signal. (**c**) UMAP showing that single cells from replicate 3 cluster according to the K562 and H1-hESCs cell types based on either the Pol2S5p peaks (left) or H3K27me3 peaks (right) deconvolved from replicate 1. (**d**) Same as (c) but showing the cells from replicate 1 plotted on the deconvolved peaks from Replicate 3.
