## Additional file 5 for "Simultaneous CUT&Tag profiling of the accessible and silenced regulome in single cells"

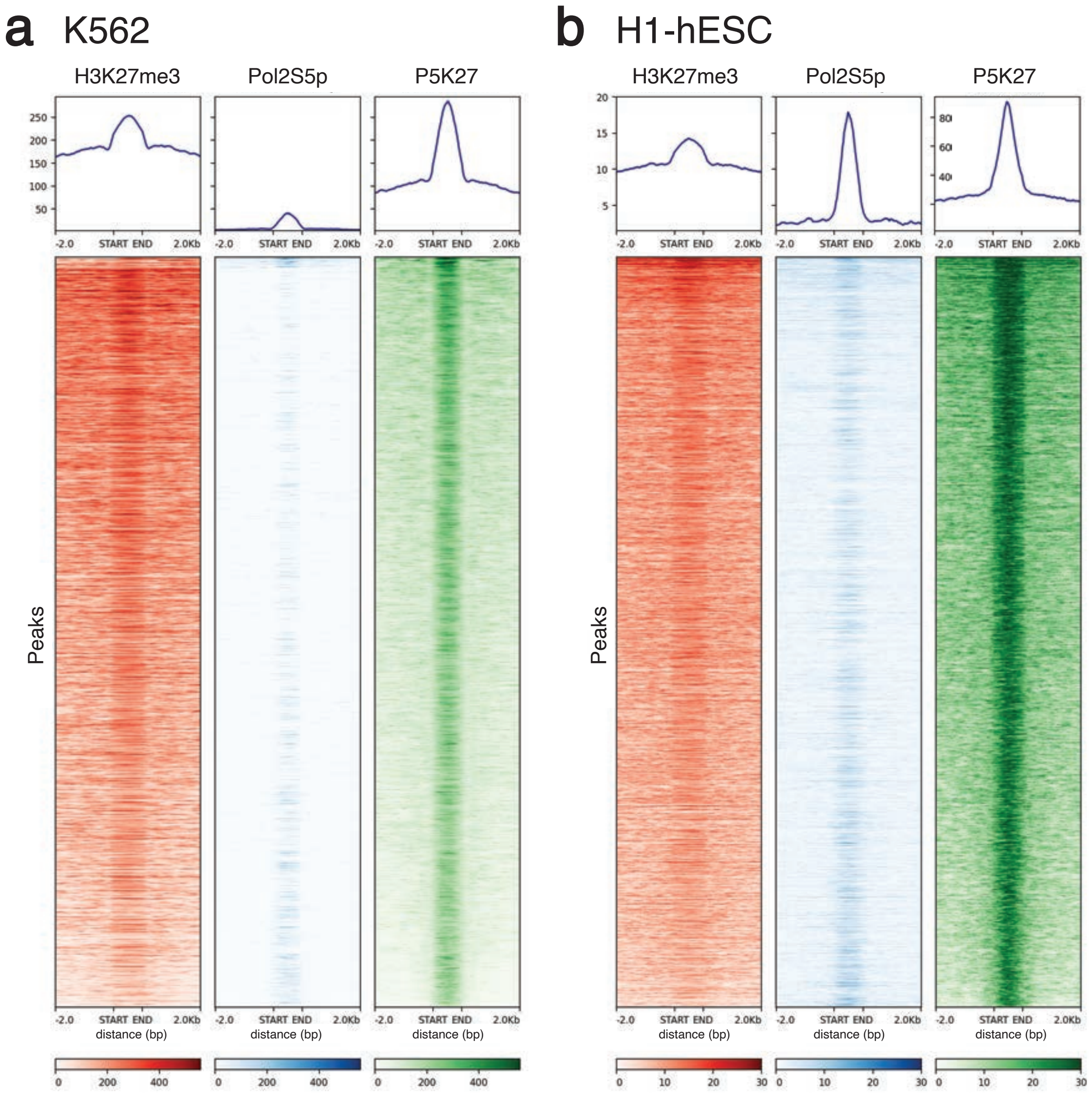


**Additional file 5:** (**a**) Single antibody and 2for1 data at the overlapping peaks for K562 cells. (**b**) Same as (a), for H1-hESC cells.
